# Detection of PCR chimeras in adaptive immune receptor repertoire sequencing using hidden Markov models

**DOI:** 10.1101/2025.02.21.638809

**Authors:** Mark Chernyshev, Aron Stålmarck, Martin Corcoran, Gunilla B. Karlsson Hedestam, Ben Murrell

## Abstract

Adaptive Immune Receptor Repertoire sequencing (AIRR-seq) has emerged as a central approach for studying T cell and B cell receptor populations, and is now an important component of studies of autoimmunity, immune responses to pathogens, vaccines, allergens, and cancers, and for antibody discovery. When amplifying the rearranged V(D)J genes encoding antigen receptors, each cycle of the Polymerase Chain Reaction (PCR) can produce spurious “chimeric” hybrids of two or more different template sequences. While the generation of chimeras is well understood in bacterial and viral sequencing, and there are dedicated tools to detect such sequences in bacterial and viral datasets, this is not the case for AIRR-seq. Further, the process that results in immune receptor sequences has domain-specific challenges, such as somatic hypermutation (SHM), and domain-specific opportunities, such as relatively well-known germline gene “reference” sequences. Here we describe CHMMAIRRa, a hidden Markov model for detecting chimeric sequences in AIRR-seq data, that specifically models SHM and incorporates germline reference sequences. We use simulations to characterize the performance of CHMMAIRRa and compare it to existing methods from other domains, we test the effect of PCR conditions on chimerism using IgM libraries generated in this study, and we apply CHMMAIRRa to four published AIRR-seq datasets to show the extent and impact of artifactual chimerism.

## INTRODUCTION

Adaptive Immune Receptor Repertoire sequencing (AIRR-seq) applies amplicon next-generation sequencing to B and T cell receptor (BCR, TCR) V(D)J transcripts. AIRR-seq is used to study receptor populations in autoimmunity, immune responses to pathogens, vaccines, allergens, and cancers, and for antibody discovery.

Polymerase chain reaction (PCR) amplification of V(D)J amplicons is unavoidable in AIRR-seq, due to the DNA input concentration required for next-generation sequencing. With each PCR cycle, incomplete extension results in partially-amplified fragments which can serve as primers in subsequent cycles. When these anneal to heterologous templates, chimeric molecules are formed (Figure 1A). Indeed, PCR chimera formation rates correlate positively with PCR cycle counts [1], [2] and template similarity [1]. Amplicon sequencing, where any two template molecules are likely to be non-identical but have substantial sequence similarity in some regions, is particularly susceptible to chimerism.

**Figure 1:**
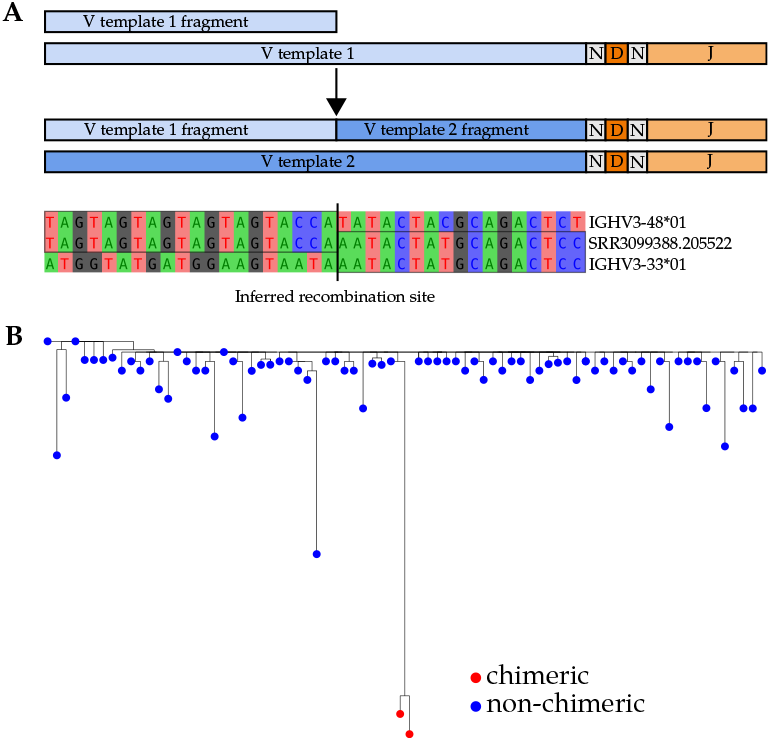
(A) Illustration of the PCR chimeric recombination process. The incompletely amplified fragment of V template 1 becomes a primer for V template 2, forming a chimera. An example chimera from a public dataset (PRJNA308641) is shown. (B) A maximum likelihood phylogenetic tree of an IgM VDJ lineage from the Chernyshev et al. 2023 dataset, depicting two chimeric sequences which resemble affinity-matured antibodies. The germline sequence was obtained from the IgBLAST-generated “germline_alignment” column of the sequence with the minimum IGHV SHM in the lineage.

Undetected PCR chimeras can confound AIRR-seq analysis. For example, during BCR lineage analysis [3], [4], [5], [6], [7], chimeric sequences can be misinterpreted as sequences with high somatic hypermutation (SHM) (Figure 1B). Besides just perturbing summary statistics, such as diversity metrics and average SHM, high SHM sequences might be of special interest and prioritized for phenotypic characterization, as SHM can correlate with antigen binding affinity. In such cases, undetected chimerism could be especially wasteful of experimental effort and resources.

There are several existing methods for chimera detection, both de novo approaches and those using reference databases. However, these methods were designed with two primary purposes in mind: the detection of PCR chimeras generated during 16S sequencing (UCHIME, ChimeraSlayer) [1], [8], and detection of real recombination events which occur during viral evolution (3SEQ, RDP5, MaxChi, Chimaera) [9], [10], [11], [12].

AIRR-seq data analysis requires consideration of its unique characteristics. The human adaptive immune receptor loci contain different numbers of variable (V), diversity (D), and joining (J) germline genes. For TCRs, individuals typically possess approximately 38 alpha specific variable (TRAV), 48 beta variable (TRBV), 8 gamma variable (TRGV), 3 delta specific variable (TRDV) genes and 5 shared TRAV/TRDV genes. For immunoglobulins, there are approximately 50 heavy chain variable (IGHV), 50 kappa variable (IGKV), and 40 lambda variable (IGLV) genes. The numbers of D and J genes show similar diversity across loci and heterozygosity is common across Vs, Ds, and Js. Though variation in these loci is high, there exist relatively complete databases for each locus and multiple methods for obtaining individual genotypes [13], [14], [15], [16]. Additionally, V(D)J sequence changes introduced by SHM can make it more difficult to reliably identify the recombined V gene. Furthermore, The frequency distribution of the sequenced repertoire, which can be methodologically relevant for detecting chimeras, can be skewed away from uniformity by clonal lineage expansion for both BCR and TCR repertoires. For B cells especially, the degree of skewing can also depend on the cell population that is being sequenced. Immunoglobulin G (IgG) isotypes exhibit more skewing due to more extensive clonal expansion than Immunoglobulin M (IgM) isotypes (which have higher proportions of unexpanded naive cells), and populations that include plasma cells exhibit skewing due to extremely high per-cell transcript counts. Despite these differences, BCR and TCR repertoires are far more uniform in distribution than those for microbial populations.

Here we propose a Chimera-detecting Hidden Markov Model for AIRR data (CHMMAIRRa) that exploits the existence of germline reference sequence databases and explicitly models SHM (for BCR queries). CHMMAIRRa attempts to explain a given query sequence as either a mutated copy of any non-chimeric reference template, or as a (possibly mutated) chimeric product of two or more reference sequences, marginalizing over all possible such explanations. The marginal probability that a query is chimeric can then be used to filter sequences, with a threshold that depends on the application.

## MATERIALS AND METHODS

### CHMMAIRRa

CHMMAIRRa detects chimeric V(D)J sequences in AIRR-seq datasets using a hidden Markov model (HMM) to model recombination as switching between reference sequences. Only the V gene, whose length spans the majority of the sequence, is considered. CHMMAIRRa takes, as input, an AIRR-formatted file, which includes the alignment of a query V(D)J against the closest matching reference sequence (generated with a standard alignment tool such as the Immunoglobulin Basic Local Alignment Search Tool (IgBLAST)), and a multiple sequence alignment (MSA) of the database V sequences. First, the query is threaded onto the reference MSA using the existing AIRR-seq alignment, exploiting the standard format to avoid having to realign against the database. Then the HMM is run over the MSA-aligned query sequence, where the terminal states in the HMM (ie. at the final V-gene alignment column) distinguish between chimeric and non-chimeric sequences (Figure 2).

**Figure 2:**
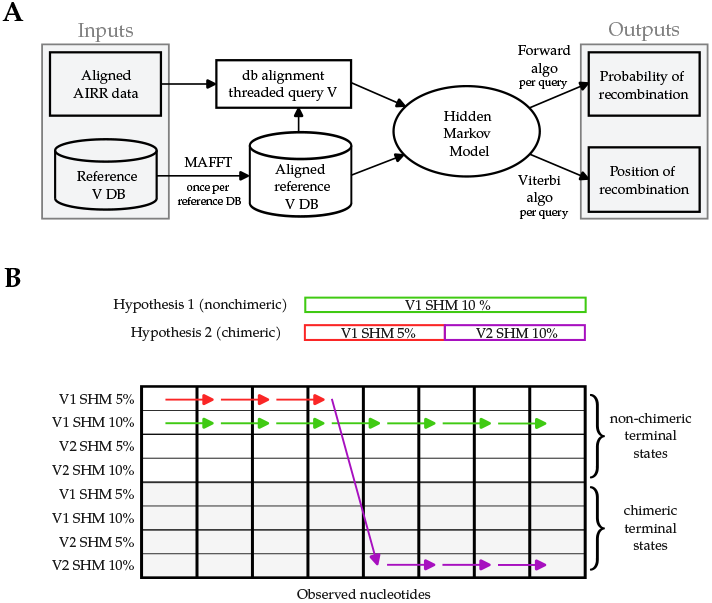
A diagram of the CHMMAIRRa pipeline. The upper half illustrates the elements of the pipeline, while the lower one depicts a an example dynamic programming lattice for the HMM. If a sequence is non-chimeric (hypothesis 1), it will remain in the same state (same V, same SHM rate) for all observations. If a sequence is chimeric (hypothesis 2), it will switch from a non-chimeric state to a chimeric one, potentially switching SHM rates.

### A Hidden Markov Model of chimeric sequences

To allow the terminal states in the HMM to distin-guish between chimeric and non-chimeric sequences, CHMMAIRRa duplicates the reference sequences in the HMM lattice, such that each state in the HMM corresponds to either a so-far “non-chimeric” state, without template switching in any prior nucleotides, or to a “chimeric” state which has switched from a different state at least once prior. Further, to allow for varying rates of SHM, and recombination between sequences with varying SHM rates, the SHM rate spectrum is discretized and the reference sequences (both non-chimeric and post-chimeric) are repeated multiple times, each with a different SHM rate. Non-self HMM transitions only allow switching into the post-chimeric state, such that any HMM path that ends in a non-chimeric state never switched at all, and any path ending in a post-chimeric state switched at least once. We use the Forward algorithm to compute the marginal probability that a query was emitted by any chimeric HMM path, and (optionally) the Viterbi algorithm to obtain the most likely parent state(s) and breakpoint locations.

For *N* germline references *R* = {*R*_1_, *R*_2_, …, *R*_*N*_}, and *K* discretized mutation rates *M* = {*M*_1_, *M*_2_, …, *M*_*K*_}, the HMM has 2*NK* states, with *NK* non-chimeric states 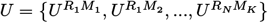 and *NK* chimeric states 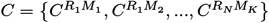, collectively *S* = {*S*_1_, …, *S*_2*NK*_}.

We model the *L* nucleotides in the observed query sequence *O* = *O*_1_, *O*_2_, …*O*_*L*_(where *O*_*t*_is the nucleotide at position *t* in the query sequence) as generated from an unobserved sequence of germline states *D* = {*d*_1_…*d*_*L*_}. If *S*_*it*_is the nucleotide of the reference sequence associated with state *S*_*i*_ when *d*_*t*_= *S*_*i*_, we set the conditional probability of the observation given the state *b*_*i,t*_(*O*_*t*_) = *P* (*O*_*t*_| *d*_*t*_= *S*_*i*_) as 1 - m if the nucleotide is the same as the reference and not a gap or N, m/3 if the nucleotide is different from the reference and not a gap or N, and 1 if the nucleotide is a gap or N.

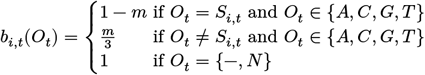

Where *S*_*i,t*_is nucleotide *t* in the reference corresponding to state *i*. The HMM does not consider biases in the reference sequence and SHM mutation probabilities, and the initial state probabilities are uniformly distributed over all *NK* non-chimeric reference states (and zero otherwise):

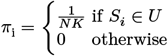

We define *R*(*i*) as the set of states corresponding to the same reference as *S*_*i*_, only including chimeric states if *S*_*i*_is chimeric, and only non-chimeric states if *S*_*i*_ is non-chimeric. We introduce two switching rate parameters, *ψ* and *μ*, where *ψ* is the rate of switching to a state corresponding to a different reference, and *μ* is the rate of switching to a state corresponding to the same reference but with a different mutation rate. We enforce the interpretation of the non-chimeric and chimeric states by setting the transition probabilities to be 1 - *ψ* – *μ* for self transitions, 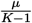 for any non-self transition into states of the same reference but with a different mutation rate, 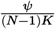 for non-self transitions into chimeric states of a different reference, and 0 for transitions into non-chimeric states of a different reference:

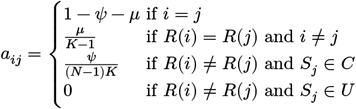

The standard HMM “Forward” algorithm (following Rabiner [17]) constructs a lattice of *α*_*t*_(*i*) = *P* (*O*_1_, *O*_2_, …*O*_*t, dt*_= *S*_*i*_) joint probabilities of the partial observation sequence *O*_1_, *O*_2_, …*O*_*t*_for all possible paths up to *t* that end in state *S*_*i*_. By construction, we can directly calculate the joint probability of observing a given query sequence and terminating in a chimeric state (which requires at least one recombination event) by summing over the second half of the final column of the lattice:

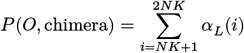

and thus, by Bayes’ theorem, we can calculate the probability that a query is chimeric:

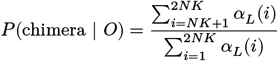

Since this handles SHM via discretizing the mutation rates, we call this approach the “Discretized Bayesian” (DB) method. We additionally considered a version of the model that has only 2 states per germline, non-chimeric and chimeric, and handles SHM by estimating a continuous per-reference mutation rate using the Baum-Welch algorithm [17], [18] (the “BW” method).

Computing HMM Forward probabilities (and reestimating parameters via Baum-Welch) using the standard algorithms is quadratic in the number of states, and here would be *𝒪*(*L* × (*NK*)^2^), which would be prohibitive for a large number of germline reference sequences. However, due to the structure of our transition probability matrix we can avoid this full pairwise computation from all states to each other for each step in the HMM lattice.

The standard induction step for the Forward algorithm is:

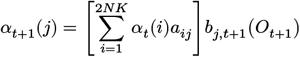

With our transition probabilities, the induction step can be rewritten as:

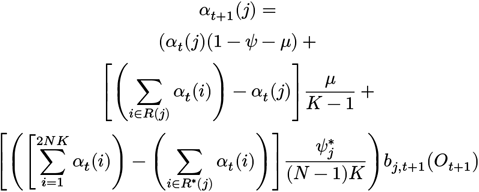

where 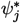 is 0 if *j* is a non-chimeric state and *ψ* if *j* is a chimeric state, and *R*\*(*j*) is the set of all states, either chimeric or non-chimeric, corresponding to the same reference. Note that 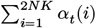 *α* (*i*) can be precomputed once and reused for each *j*, and ∑_*i*∈*R*(*j*)_ *αt*(*i*) has a complexity of *𝒪*(*K*) and can be precomputed N times and reused for each mutation rate, reducing the total complexity to linear in the number of HMM states: *𝒪*(*LNK*)

The same approach applies to the backwards algorithm, while the Viterbi algorithm requires a precomputation of a maximum instead of a sum.

### IgM library preparation

24 5′ multiplex IGM libraries were prepared based on the protocol previously described [19]. Libraries were prepared with varying starting template total RNA and Indexing PCR template, modifications in the number of PCR cycles in the amplification step, and changes to the length of the extension step. See supplementary data 1 for the full list of conditions.

### AIRR-seq data preprocessing

In this study, we analyzed four published datasets and one new dataset. The new dataset is composed of 24 IgM repertoire libraries designed to study the effects of PCR protocol variation on chimerism. For all libraries, we inferred the set of germline alleles, and annotated the sequences, using IgDiscover22 [13] v1.0.0. Detailed pre-processing settings and instructions are included in the paper’s GitHub repository (see code availability section).

Dataset-specific preprocessing:

- Chernyshev et al [7] and Corcoran et al [20]: as described in their respective publications.
- Rubelt et al twins dataset [21] (PRJNA300878): fastq files were combined per individual then split by TRA/TRB constant region presence. IgDiscover was run with TRA/TRB specific settings.
- Galson et al. [22] (PRJNA308641): database and repertoire sequences were trimmed to account for IGHV region primers, then IgDiscover was run with barcode length 0. Personalized germline IGH VDJ sets identified by running IgDiscover on IgM libraries were used to assign IgG sequences from the same donor.
- This paper’s dataset: IgDiscover was run on control (condition 1/C1) libraries to determine personalized IGH VDJ sets, which were then used to assign sequences belonging to the corresponding donor.

### Comparison methods

While not designed for this specific immunoglobulin repertoire sequencing use-case, some existing tools can identify chimerism and we compared our approach to:

1. USEARCH: We used USEARCH v11.0.667_i86linux32. Only results marked with the ‘Y’ annotation were interpreted as chimeras.

2. VSEARCH: We used the VSEARCH v2.18.0_linux_x86_64 uchime_ref command with the default score threshold of 0.28.

All methods were run on 16 threads using an Intel(R) Xeon(R) Gold 6134 CPU @ 3.20GHz.

## RESULTS

### Simulated data

To evaluate method performance, we simulated six datasets: two using a synthetic TRBV genotype constructed by randomly (uniform) selecting one allele per gene from the KI human TRBV database (release v0.0.1) [20] and four using a synthetic human IGHV genotype constructed by randomly (uniform) selecting one functional allele per gene from the Open Germline Receptor Database (OGRDB) Homo sapiens IGHV germline set (release 9) [23]. Each dataset contained 10,000 sequences with a 5% chimerism rate (500 recombined, 9,500 non-recombined) and was simulated at varied percent differences from reference (DFR) (0.5%, 2% for TRB and 0%, 5%, 10%, and 20% for IGH). DFR represents sequencing errors in TCRs and SHM in BCRs. Non-recombined sequences were generated by uniformly mutating a single reference sequence, while recombined sequences were created by combining two uniformly mutated references at a breakpoint along the sequence selected uniformly and at random.

### CHMMAIRRa Parameters

For simulated data, which aims to compare our two approaches to each other, and to previous methods, we ran CHMMAIRRa in three configurations: the DB approach with the mutation rate set to 0.005 for TRB data, the DB approach with the SHM rate discretized into 15 categories (from 0.0 to 0.25) for IGH data, and the BW approach which reestimates the mutation rate (initialized at 0.05). For real data analysis, we used CHMMAIRRa’s --receptor parameter, which automatically selects appropriate settings for each receptor type: for immunoglobulin data we use BW mode (initialized to a 0.05 mutation rate), and for TCR data, which lacks SHM, we use DB mode with a fixed 0.005 mutation rate to allow for sequencing error. Full parameter settings are detailed in Supplementary Data 1.

### Performance

Figure 3A and Figure 3B show that CHMMAIRRa achieved the highest area under the receiver-operating characteristic (ROC) curve (AUC) across all tested methods when evaluating simulated TRBV and IGHV repertoires with varying DFR, with both CHMMAIRRa approaches (BW and DB) performing similarly well. CHMMAIRRa’s default detection threshold (of 0.95 posterior probability) seems appropriately calibrated to identify most chimeras, but avoid discarding useful data through false chimera detections, with zero false positives at IGHV SHM rates up to 10%, and only extremely low false positives at IGHV SHM rates of 20% (0 for DB, and 6/9500 for BW). The USEARCH uchime2_ref “balanced” threshold was similarly stable across simulation scenarios, but was dominated in call cases by CHMMAIRRa. The standard VSEARCH uchime_ref threshold performed quite poorly as IGHV SHM increased, for example detecting just over 10% of the true chimeras at 20% SHM rates.

**Figure 3:**
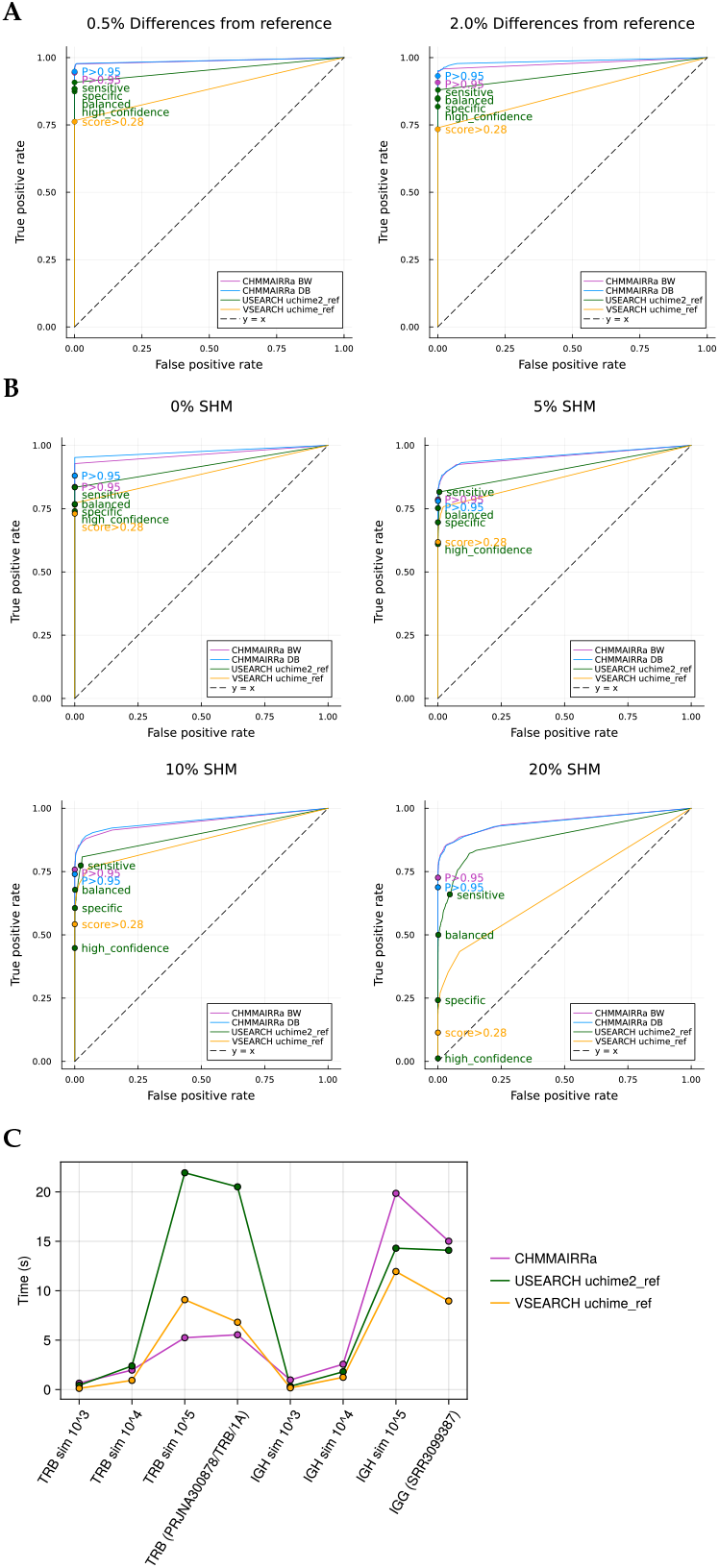
(A) ROCs depicting the performance of the chimera detection tools on simulated TRB datasets with 0.5% and 2% differences from reference (mimicking low and high sequencing error rates). (B) ROCs depicting the performance of the chimera detection tools on simulated IGH datasets with 0%, 5%, 10%, and 20% SHM rates. Default cutoffs for all methods are labelled on the ROCs. (B) Method run times on simulated and real TRB and IGH repertoire data. Simulated data was composed of 100% chimeras with a 0% mutation rate. CHMMAIRRa was run with default settings for the corresponding receptor, USEARCH with balanced mode, and VSEARCH with default settings.

Runtime analysis (Figure 3B) showed that CHMMAIRRa was competitive with existing tools, though its performance varied by receptor type. CHMMAIRRa was fastest across larger TRB datasets and slightly slower for large IGH datasets, likely because the IGH datasets require BW re-estimation of SHM rates. Note that USEARCH and VSEARCH are solving a very different computational problem, including sequence alignment to reference databases, but CHMMAIRRa exploits the fact that AIRR-seq processing pipelines already align queries to reference databases (using approaches tailored for IG data), and avoids realignment. Importantly, all chimera detection methods added negligible processing time compared to standard AIRR pipeline steps like VDJ annotation with IgBLAST.

### Reference database completeness

We hypothesized that missing databases alleles (a common scenario when personalized germline sequences are unavailable) might cause false chimera identification, because some genuine alleles can appear as recombinations between other alleles in the reduced database, and investigated CHMMAIRRa’s robustness to incomplete reference databases. Using both IGH and TCR data, we compared chimera detection performance across complete V gene databases and random (uniform) subsampled versions. Full database details are provided in Supplementary data 1.

For IGH:

- OGRDB [23]
- IMGT [24]
- IgDiscover inferred personalized genotypes (PG) [13]
- Random PG subsets (30 or 40 Vs)

For TCR:

- KI TCR database [20]
- IMGT [24]
- IgDiscover PGs
- Random PG subsets:
- TRA: 30 or 40 Vs
- TRB: 40 or 50 Vs
- TRG: 4 or 6 Vs

Confirming our hypothesis, random database subsampling led to inflated chimera detection rates (Figure 4). Detected chimerism is, however, relatively consistent between PG databases, which contain novel alleles specific to these datasets, and two published non-personalized databases, suggesting that public, non-personalized, databases can indeed be used for chimera detection, as long as they are sufficiently complete.

**Figure 4:**
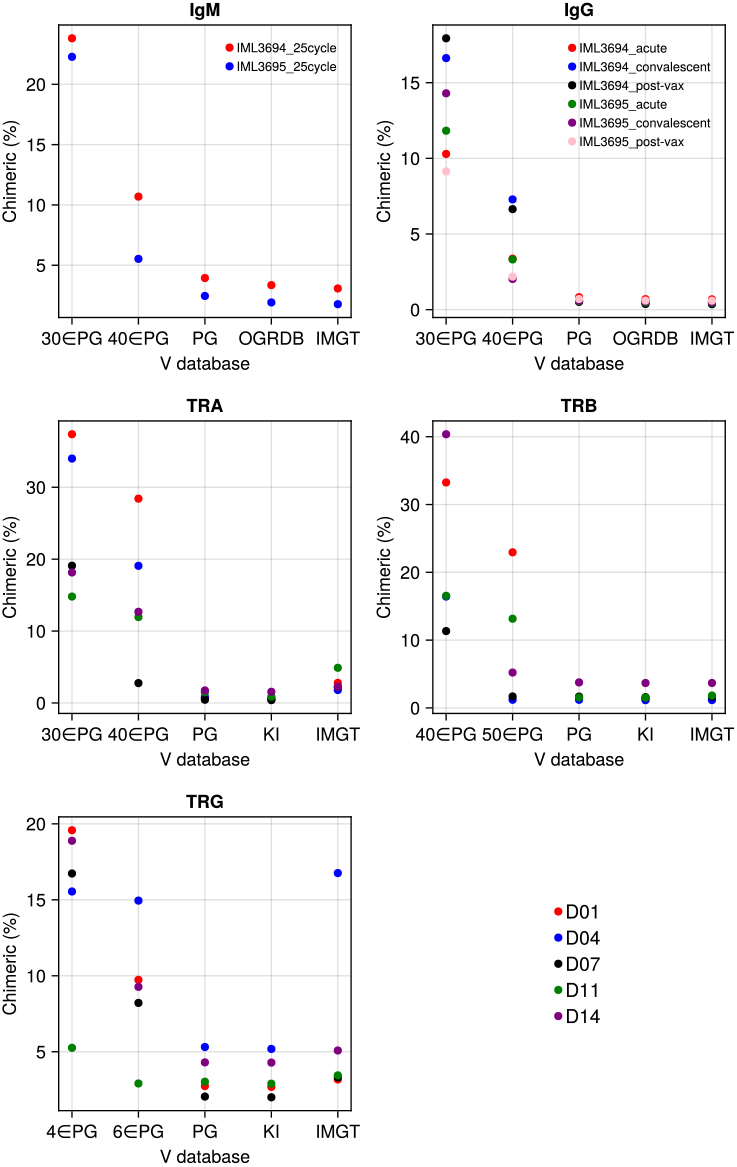
Personalized genotypes, the KI TCR database, and IMGT database all yield consistent chimerism rates. Subsampling the personalized genotype results in a sharp increase of chimera detection. Methods settings listed in Supplementary data 1.

TCR data is more susceptible to misidentifying missing alleles as chimeras due to its low tolerance for mutations (as the model only expects sequencing errors). In Figure 4, library TRG/D04 from the Corcoran et al. dataset shows elevated chimerism when analyzed with the IMGT database because it contains a highly expressed allele (TRGV4*02_S0072) that is absent from IMGT. To help identify such cases, CHMMAIRRa provides a visualization of normalized recombination counts (Supplementary Figure 1), where missing alleles appear as overrepresented recombinations when comparing analyses using databases with and without a missing allele.

Our analysis of the Corcoran et al. TCR dataset [20] revealed an interesting case where CHMMAIRRa consistently flagged TRGV7*01 as a TRGV8*01/TRGV3*01 recombinant across all donors. While IMGT annotates TRGV7*01 as a pseudogene, it contains a functional recombination signal in the GRCh38 assembly and was found in all 45 TRG libraries. This annotation discrepancy has led to inconsistent treatment in the literature:

Gherardin et al. [25] reported substantial TRGV7 expression, while Mimitou et al. [26] excluded it from their analyses. This shows that chimera detection can also uncover other technical issues for rep-seq analysis, if aberrant chimerism patterns are appropriately investigated.

### Effects of PCR protocol variation on detected chimerism

To confirm our understanding about the experimental origins of the chimeras we aim to detect, we investigated how PCR protocol variations affect chimera formation by generating IgM VDJ repertoire libraries under eight different conditions, varying:

- PCR cycles (25, 30, 35, 45)
- Indexing PCR template (5, 50 ng)
- Input RNA (25, 250, 1000 ng)
- Extension time (1, 30 seconds)

Consistent with previous studies from other biological systems [2], [27], [28], both PCR cycle number and template concentration showed strong positive correlations with chimera formation (Figure 5). Chimerism rates began to plateau at 45 cycles, possibly because dNTP or primer depletion made any further cycling unproductive. Input RNA amount had minimal impact across the tested range (250-1000 ng). Unlike Qiu et al. [27], we found no effect of extension time, suggesting that 1 second is sufficient for our amplicon length when using the KAPA HiFi Hotstart ReadyMix system.

**Figure 5:**
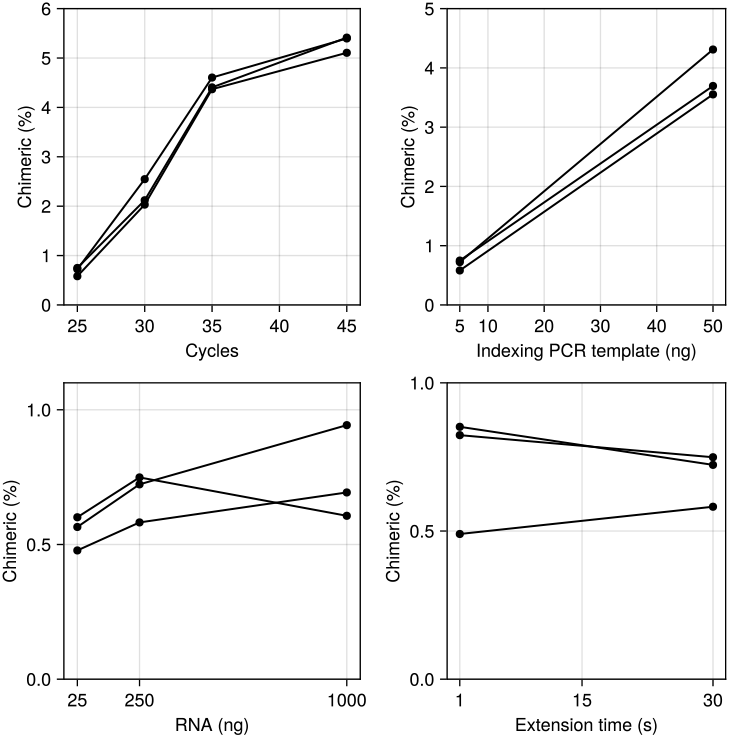
The effect of PCR cycles, Indexing PCR template, RNA input, and PCR extension time on chimera rate. The number of cycles and indexing PCR template are positively correlated with chimerism rates. See Supplementary data 1 for the full list of PCR conditions tested. Methods settings listed in Supplementary data 1.

### Detected chimerism in human AIRR-seq data

We used CHMMAIRRa to analyze chimera formation patterns across 319 libraries from four datasets published by 3 labs spanning all adaptive immune receptor chains (Figure 6). Chain-specific differences in chimera formation were evident: TRG and IGH libraries showed the highest rates, while TRD libraries contained almost no chimeras. These patterns correlate with the genetic distances between germline alleles within each chain. Specifically, genes separated by edit distances of ∼10-80 nucleotides formed more chimeric recombinations than expected based on their individual frequencies (Supplementary Figure 2), compared to more diverged genes, or nearly identical genes. This is expected as some homology is needed for the chimera formation process, and some divergence is required for chimera detection. The near-absence of chimeras in TRD libraries likely reflects the unique structure of the TRDV germline set, which lacks alleles within this critical distance range (Supplementary Figure 3, Figure 6).

**Figure 6:**
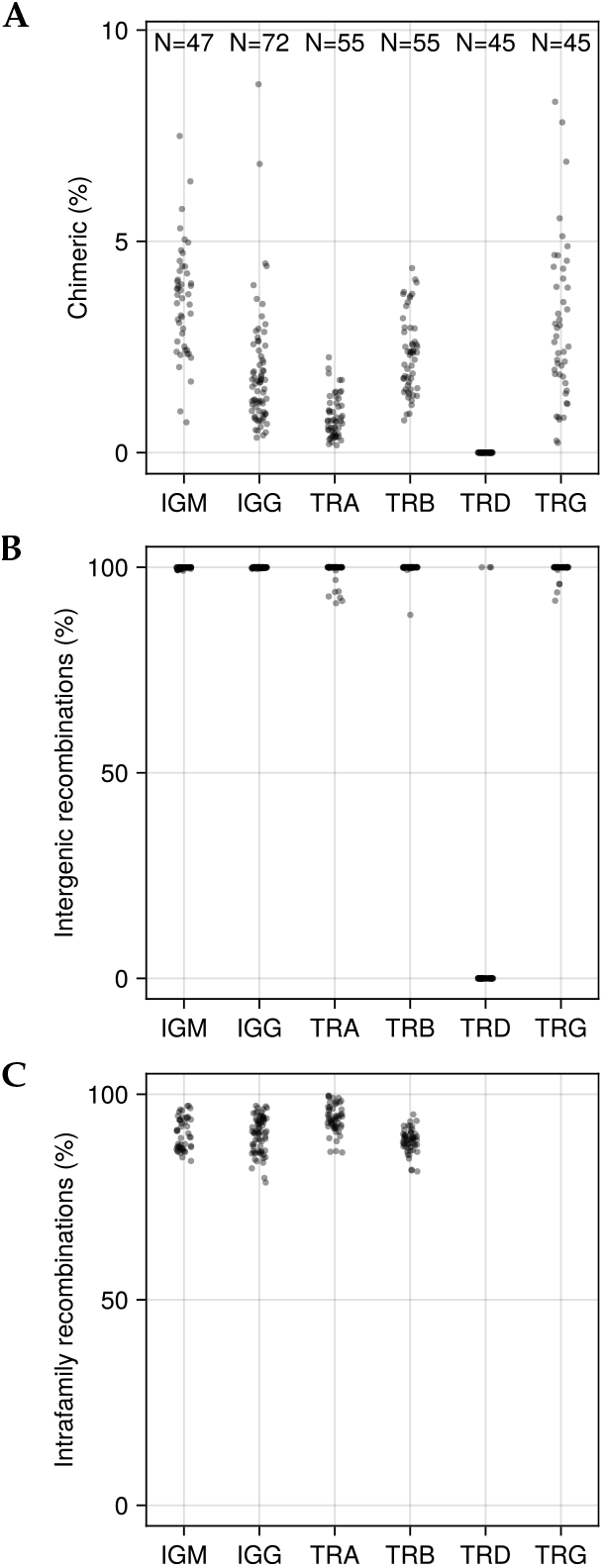
Scatter plots of (A) Percent chimeric sequences per library. (B) Percent of chimeras with fragments from different genes. (C) Percent of chimeras with fragments from the same families. TRD and TRG loci were excluded since they do not have gene families, only genes. Methods settings listed in Supplementary data 1.

Within BCR datasets, IgM libraries consistently showed higher chimerism rates than paired IgG libraries from the same individuals. While this pattern was reproducible across datasets, it could reflect either genuine differences in PCR chimera formation or reduced detection sensitivity in the more heavily mutated IgG sequences. Most detected chimeras occurred between V genes within the same family (Figure 6), confirming that sequence similarity is critical for chimera formation.

The patterns of gene combinations in chimeric sequences were similar between IgM and IgG libraries. In the TRGV locus, we observed notably low chimerism for three genes (Figure 7): TRGV5P and TRGV1, likely due to their low expression levels, and TRGV9, due to its high sequence divergence from other TRGV genes.

**Figure 7:**
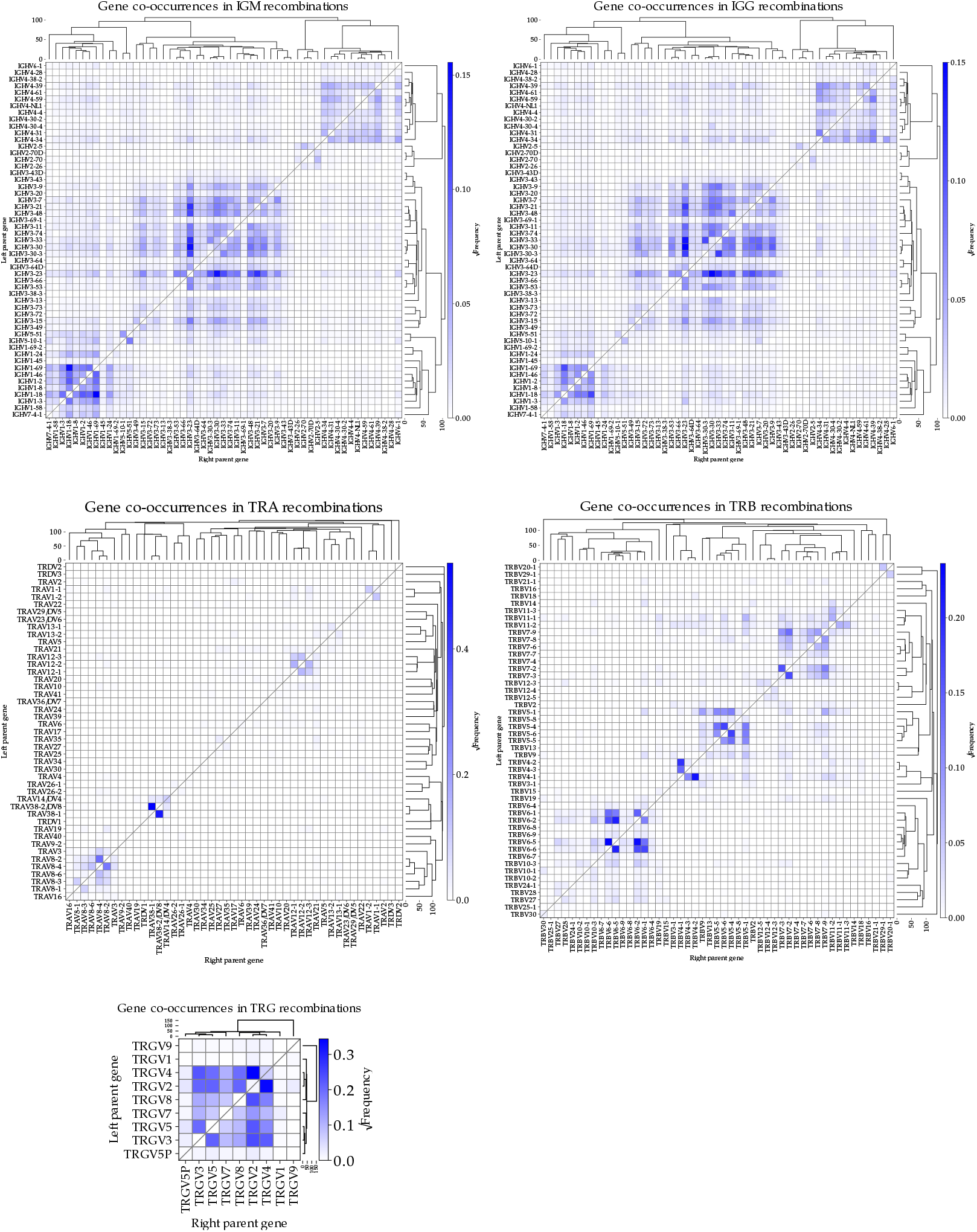
Heat maps depicting the square root of the frequency of gene to gene recombinations detected by CHMMAIRRa across all (A) IgM libraries (other than the PCR conditions tests), (B) IgG libraries, (C) TRA libraries, (D) TRB libraries, and (E) TRG libraries. The rows of the heat maps are genes on the left side of the recombined sequence, while the columns are genes on the right side. The dendrograms represent average linkage hierarchical clustering of a pairwise distance matrix of average gene to gene distances. Only single recombination chimeras were included. Methods settings listed in Supplementary data 1.

## Discussion

We present CHMMAIRRa, a hidden Markov model-based tool for detecting chimeric sequences in AIRR-seq data. CHMMAIRRa is tailored specifically for the AIRR-seq setting, where there is i) an exploitable reference sequence database, ii) the sequences can be relatively well aligned, and iii) some of the sequences can be extensively mutated. While on unmutated sequences its performance is similar to existing tools developed for other settings (USEARCH uchime2_ref and VSEARCH uchime_ref), CHMMAIRRa shows superior accuracy when analyzing sequences with high somatic hypermutation. This improved discrimination between true mutations and chimeric artifacts is critical for AIRR-seq applications because chimeric sequences can:

1. Mimic highly affinity-matured antibodies, potentially misdirecting therapeutic antibody discovery
2. Inflate diversity metrics in immune response studies
3. Interfere with linage clustering
4. Distort evolutionary analyses by introducing artificial mutation patterns

“Perfect fake chimeras” are genuine sequences that are indistinguishable from combinations of two other sequences [8], and can present a challenge for chimera detection in some settings. CHMMAIRRa addresses this challenge through its probabilistic framework, which evaluates each sequence as either a mutated single template or a combination of mutated templates. This approach naturally expresses uncertainty when analyzing potential perfect chimeras with similar parent sequences. However, when parent sequences are more divergent, the likelihood of a genuine sequence perfectly matching their combination decreases, leading to higher confidence in chimera detection. Users can fine-tune this trade-off by adjusting CHMMAIRRa’s probability threshold and prior recombination probability parameters.

While neither USEARCH uchime2_ref nor VSEARCH uchime_ref were specifically designed to handle the high mutation rates characteristic of SHM, USEARCH uchime2_ref performs better in these high-noise scenarios, possibly due to its alignment strategy and possibly due to its scoring system. We attribute CHMMAIRRa’s performance to its probabilistic formulation that considers all possible non-chimeric and chimeric explanations for a given query, weighted by their respective probabilities under the model, explicitly allowing SHM.

Our analysis of the Corcoran et al. TRG libraries yielded an unexpected insight: sequences initially flagged as TRGV8*01/TRGV3*01 chimeras were actually TRGV7*01, a gene annotated as a pseudogene in IMGT. This gene differs from its closest functional relative (TRGV3*01) by 35 nucleotide changes and a deletion. Such large differences pose challenges for current germline inference methods, which typically focus on small variations from reference sequences and often ignore insertions/deletions [13], [15]. This case, as well as the TRGV4*02_S0072 allele in donor D04, demonstrate how chimera detection tools can help identify missing or misannotated genes, hinting at a broader utility of these methods. In addition to the core chimera detection functionality, our package includes visualization tools that assist in identifying such cases.

Detected AIRR-seq library chimerism rates can be influenced by PCR protocol parameters (including cycles and indexing PCR template), the properties of the receptor’s germline V set (such as the presence of highly expressed and similar V alleles), as well as the completeness of available germline databases. For our specific amplification and library preparation protocol, [19] 5 ng of indexing PCR template material, 25ng of RNA input and 25 PCR cycles provides a good balance between minimizing chimerism and ensuring detection of rare sequences in IgM libraries. Across libraries, chimera formation rates vary substantially from one chain to another, the most extreme being the TRDVs with nearly zero chimeras, as well as IgMs and TRGVs with the highest levels of chimerism. These observations are consistent with the presence of highly expressed and similar V alleles in the germline repertoires. For AIRR-seq, excluding the two TRGV examples discussed, detected chimerism did not vary between complete databases and personalized genotypes. Therefore, while personalized genotypes are always preferred when available, in their absence a relatively-complete reference databases suffices for accurate chimera detection. Our investigation here was with human data, and database completeness varies by species, so if used for other species the effects of missing reference should be explicitly investigated.

While standard HMM algorithms are quadratic in the number of states, we exploit the structure of our transition matrix to reduce this to linear, which was critical for efficiency when the number of reference sequences is large. The assumptions required for this may apply to many other situations, and while we are not sure whether our approach is completely novel (as the HMM literature spans many fields and many decades), there are certainly related domains where this could be applied, such as identifying real (ie. not PCR induced) recombination events in biological sequences, where some tools already rely on HMMs [10]. Our implementation is sufficiently modular to allow the core linear algorithm to be reused for these other contexts.

## Limitations

CHMMAIRRa has some limitations. Primarily, detection is limited to the V region, excluding D and J regions. We are not optimistic about identifying chimeras with breakpoints in the D region, but it may be possible to identify J region chimeras, if they are between suitably diverged J alleles. We consider this a problem for future work. Also, as CHMMAIRRa is reference-based, it is unable to identify chimerism between two different templates from the same V allele. There are some non-reference based strategies that might be able to tackle this problem in some cases, such as recombination between high frequency clonal lineages with distinguishing mutations, but we expect that such cases are likely not sufficiently common for this to justify the additional computational expense.

## Conclusion

AIRR-seq is becoming a central tool across immunology and in various biotechnology sectors, and the areas where immune repertoires are considered relevant is rapidly expanding. Here we describe PCR chimera formation as a key source of error in these datasets, and release a computationally inexpensive tool to ameliorate it. We argue that any AIRR-seq processing pipeline should aim to quantify and remove PCR chimeras, both during experimental protocol development (so that conditions can be optimized to reduce chimera formation) and during routine repertoire sequence analysis.

## Data availability

- This paper’s IgM repertoire libraries generated to test the effect of PCR protocol variation on chimerism are available on the European Genome Archive under the accession number EGADXXXXXXXXXXX.
- Rubelt et al. [21] TRA/TRB repertoire data are available on SRA at the accession PRJNA300878.
- Galson et al. [22] IgM/IgG repertoire data are available on SRA at the accession PRJNA308641.
- Corcoran et al. [20] TRA/TRB/TRD/TRG repertoire data are available upon request from the SciLifeLab Data Repository at https://doi.org/10.17044/scilifelab.14579091.
- Chernyshev et al. [7] IgM/IgG repertoire data are available upon request from the SciLifeLab Data Repository at https://doi.org/10.17044/scilifelab.21518142.

## Code availability

CHMMAIRRa and CHMMera (containing the core HMM implementation) are published on the Julia package registry and are available at github.com/Mur-rellGroup/CHMMAIRRa.jl and github.com/Murrell-Group/CHMMera.jl. Data preprocessing and plotting scripts for this paper are at github.com/MurrellGroup/CHMMAIRRaAnalyses.

## Contributions

Conceptualization, B.M; Formal Analysis, M.C., A.S., and M.Co.; Investigation, M.C. and A.S.; Visualization, M.C. and A.S.; Writing – Original Draft, M.C., A.S., and B.M; Writing – Review & Editing, M.C., A.S., G.B.K.H. and B.M; Funding Acquisition, G.B.K.H. and B.M.

## Ethics statement

Peripheral blood mononuclear cells were sampled from three donors with informed consent under ethical permits #2016/1053-31, #2017/852-32, and #2019-01014. These samples were used to generate the 24 libraries with varied PCR conditions. All other sequencing libraries used in this study were from publicly available datasets; see data availability section.

## Supplementary figures

**Supplementary Figure 1:**
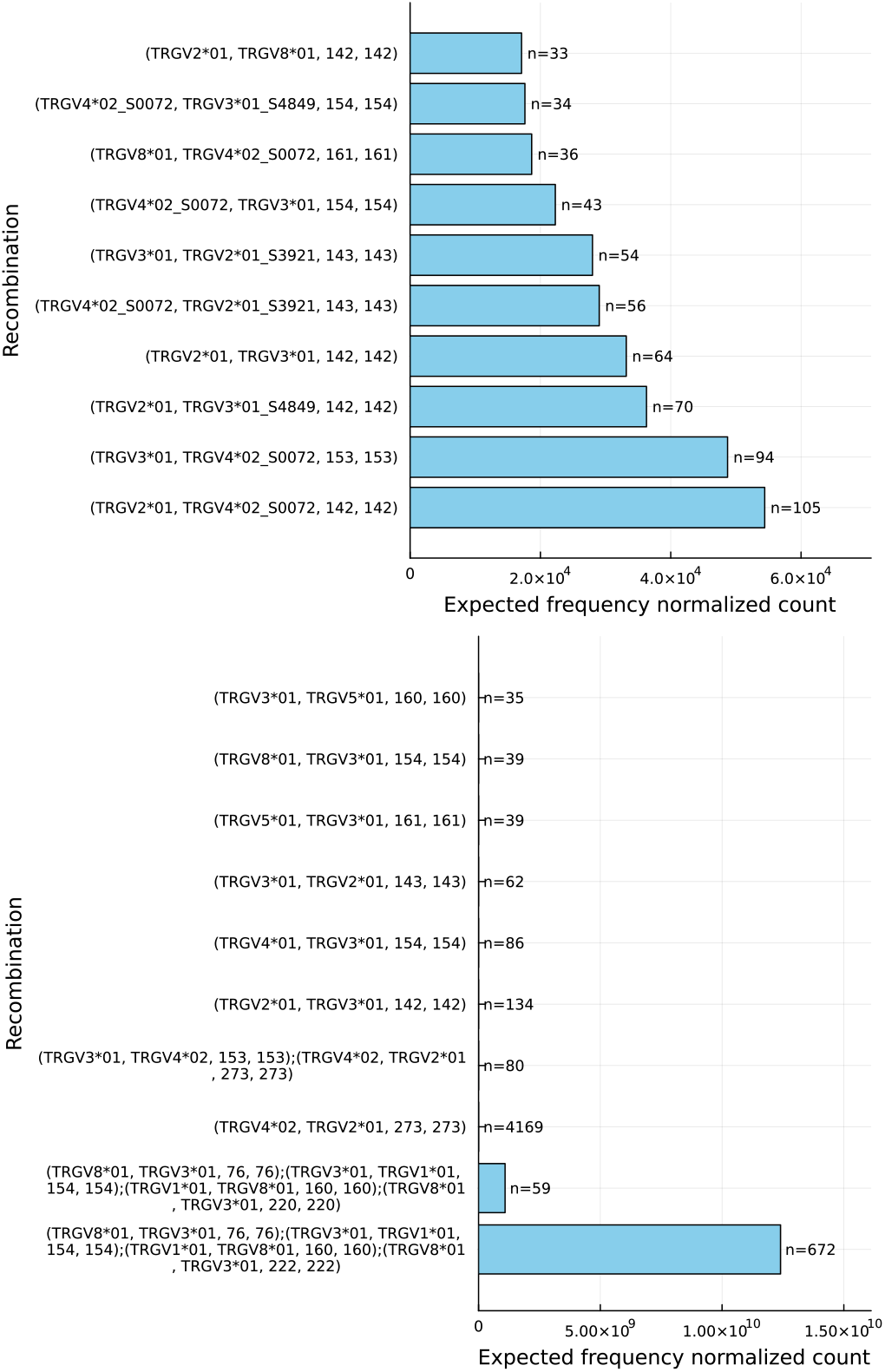
The bar plots depict the normalized counts of recombinations found by CHMMAIRRa in the TRG/D04 library from the Corcoran et al. dataset analyzed with the (A) the IMGT database and (B) the donor’s personalized genotype. Normalization was performed by dividing the occurrences of a recombination by the product of template allele frequencies. The overrepresented recombination in (B) is actually the TRGV4*02_S0072 allele, missing from the IMGT database.

**Supplementary Figure 2:**
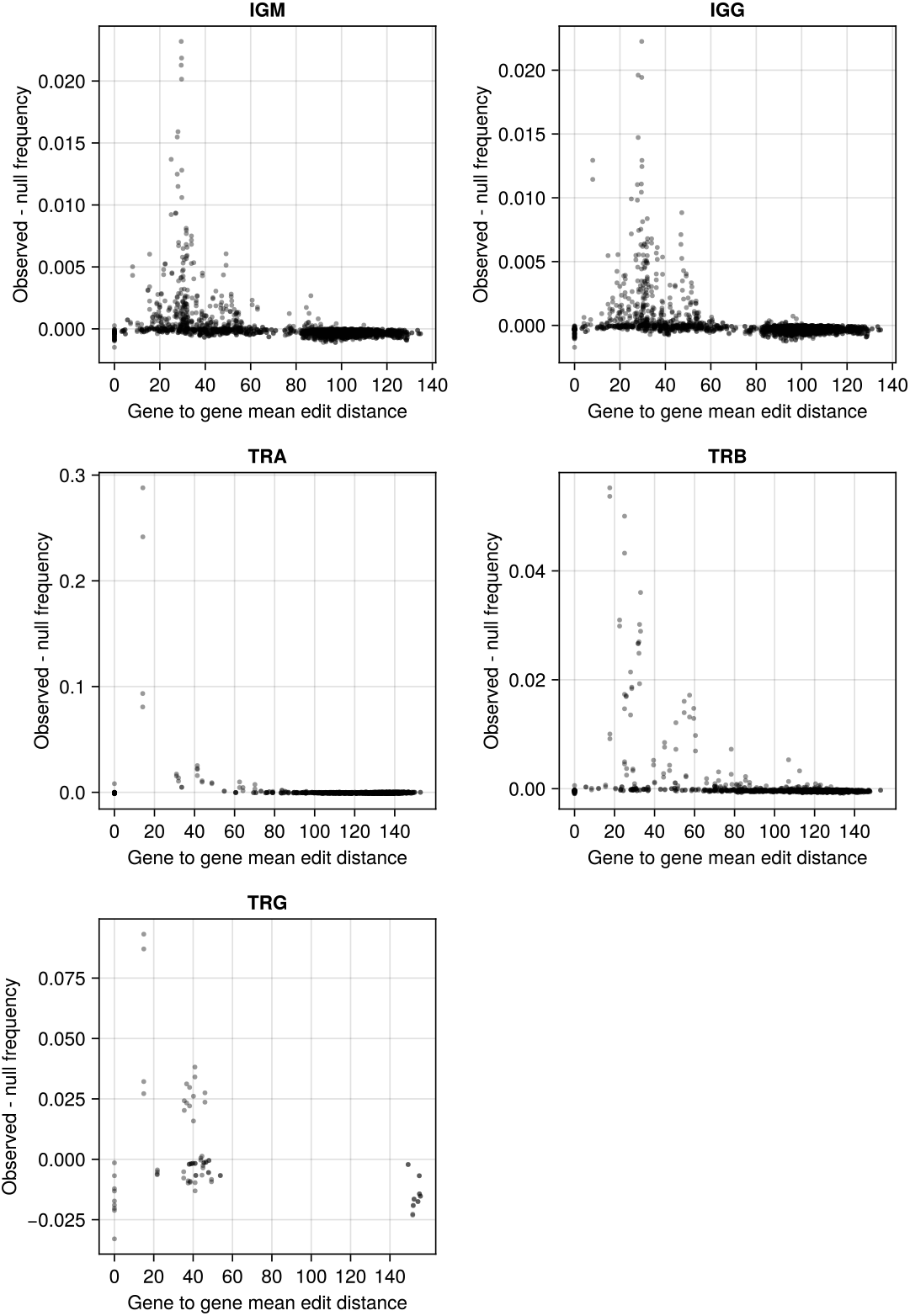
Scatterplots demonstrating the relationship between gene to gene edit distance and overrepresented gene to gene recombinations. Each dot represents all recombinations involving two specific genes. The observed frequency refers to co-occurrence of genes in recombinations, averaged across all datasets utilized in this paper except the PCR conditions test dataset. The expected frequency was calculated by taking the product of the two genes’ frequencies. Dots above y = 0 are recombinations which are observed more than expected by taking the product of the two genes’ frequencies, while dots below y = 0 are observed less often than expected. Methods settings listed in Supplementary data 1 and datasets in the data availability section.

**Supplementary Figure 3:**
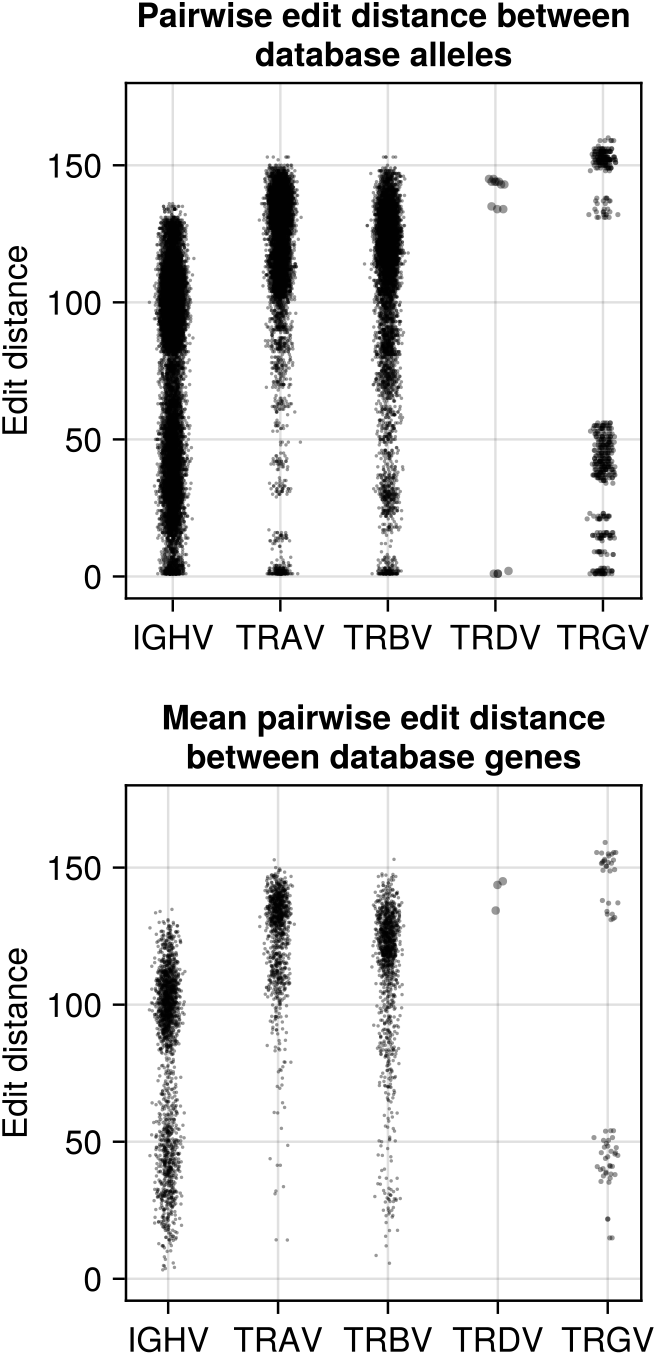
Scatterplots depicting the distribution of Levenshtein distances (A) between all pairs of V alleles and (B) averaged per-gene. The lack of similar TRDV alleles may be related to the lack of chimeras in TRD libraries. The OGRDB release 9 was used for the IGHVs and the KI TCR database v0.0.1 was used for the TCRVs. Database versions listed in Supplementary data 1.

